# Inference of direction, diversity, and frequency of HIV-1 transmission using approximate Bayesian computation

**DOI:** 10.1101/071050

**Authors:** Ethan O. Romero-Severson, Ingo Bulla, Nick Hengartner, Inês Bártolo, Ana Abecasis, José M. Azevedo-Pereira, Nuno Taveira, Thomas Leitner

## Abstract

Diversity of the founding population of Human Immunodeficiency Virus Type 1 (HIV-1) transmissions raises many important biological, clinical, and epidemiological issues. In up to 40% of sexual infections there is clear evidence for multiple founding variants, which can influence the efficacy of putative prevention methods and the reconstruction of epidemiologic histories. To measure the diversity of the founding population and to compute the probability of alternative transmission scenarios, while explicitly taking phylogenetic uncertainty into account, we created an Approximate Bayesian Computation (ABC) method based on a set of statistics measuring phylogenetic topology, branch lengths, and genetic diversity. We applied our method to a heterosexual transmission pair showing a complex paraphyletic-polyphyletic donor-recipient phylogenetic topology. We found evidence identifying the donor that was consistent with the known facts of the case (Bayes factor >20). We also found that while the evidence for ongoing transmission between the pair was as good or better than the singular transmission event model, it was only viable when the rate of ongoing transmission was implausibly high (~1/day). We concluded that the singular transmission model, which was able to estimate the diversity of the founding population (mean 7% substitutions/site), was more biologically plausible. Our study provides a formal inference framework to investigate HIV-1 direction, diversity, and frequency of transmission. The ability to measure the diversity of founding populations in both simple and complex transmission situations is essential to understanding the relationship between the phylogeny and epidemiology of HIV-1 as well as in efforts developing new prevention technologies.

## INTRODUCTION

Most HIV-1 infections are the result of sexual transmission (SHATTOCK AND MOORE 2003), where 20-40% involve transmission of multiple genetic variants (KEELE *et al*. 2008; SALAZAR-GONZALEZ *et al*. 2009; LI *et al*. 2010; RIEDER *et al*. 2011). Transmitting more than one variant raises many important biological, clinical, and epidemiological issues. Biologically, successful transmission of >1 variant means that many viruses in a donor have the capacity to establish infection, and further that they had similar fitness as they did not outcompete each other in the new host. Following establishment of infection, the existence of multiple lineages may also generate virus with higher relative fitness than when single lineages establish infection (CARRILLO *et al*. 2007), due either to recombination or competition after transmission (SANBORN *et al*. 2015). Clinically, transmission of several virus variants may make it harder for the immune system to combat the virus (GROBLER *et al*. 2004; YANG *et al*. 2005; SMITH *et al*. 2006), easier for the virus to evade antiviral treatment (SMITH *et al*. 2004), and may accelerate disease progression (GOTTLIEB *et al*. 2004). Epidemiologically, the establishment of >1 genetic variant can occur simultaneously at one time or sequentially over a long period of time, which is defined as coinfection or super-infection, respectively (VAN DER KUYL AND CORNELISSEN 2007). This has further impact on whether one infection protects against another (ALTFELD *et al*. 2002; RONEN *et al*. 2013), or if later super-infections may induce drug resistance (SMITH *et al*. 2005), and if a potential vaccine to one form would protect against another.

Phylogenetics reconstructs evolutionary history, and for an organism like HIV-1 that evolves very rapidly, the joint pathogen phylogeny from hosts that have infected each other reveals details about the host-to-host transmission. Recently, coalescent-based simulations showed that the resulting phylogeny may reveal both direction and directness in epidemiologically linked hosts, i.e., who infected whom, and whether missing host-links were likely (ROMERO-SEVERSON *et al*. 2016). Furthermore, it has previously been shown that there exists a pretransmission interval that describes the bias towards the past when using phylogenetic trees to estimate transmission times (LEITNER AND ALBERT 1999; LEITNER AND FITCH 1999; ROMERO-SEVERSON *et al*. 2014). Importantly, when multiple phylogenetic lineages have been transmitted from one host to another the resulting tree opens up alternative interpretations of whether all lineages were transmitted at one or several occasions. Thus, while simulations have shown that phylogenies carry detailed information about who infected whom, and within-host models predict the pretransmission interval, a single framework to determine the evidence for the various possible transmission scenarios between two infected hosts is lacking.

The objective of this study was to create a unified framework to investigate the nature of an epidemiological link and to apply that to a real HIV-1 transmission case. Based on previous theoretical work the tree topology should probabilistically indicate direction and directness, whether >1 lineage were transmitted, as well as when transmission occurred. Here, we also intended to determine the evidence for whether the infection was established by a single transmission event or an ongoing process of re-infection. In addition, we wanted to avoid basing our inferences on a single (best) phylogenetic tree as many trees with different topology and distance properties may be nearly as likely as the best tree. Basing our method on the entire posterior distribution of trees allows us to consider the full range of solutions that the data may support and to propagate uncertainty in phylogenetic reconstruction onto the parameter estimates. Thus, we extended our previous within-host coalescent methods to simulate trees corresponding to different transmission scenarios and parameterizations and analyzed a previously unpublished HIV-1 transmission chain. To test and compare alternative scenarios of the epidemiological link, i.e., when and how transmission(s) occurred, we developed and applied an approximate Bayesian computation (ABC) method based on tree topology, root host-assignment, and patristic tree distance measures. The ABC method also allowed us to estimate the diversity at the time of transmission rather than at time of sampling.

## MATERIALS AND METHODS

### Joint linear within-hosts population model

We considered two alternative sexual transmission scenarios: i) a singular transmission event, or ii) multiple transmissions where the donor and recipient are repeatedly re-infecting each other (Fig 1). In the singular transmission scenario the within-host effective population size, *N(t) = α + βt*, is a linear function of time where *α* is the population size at the time of infection at time *t* = 0 and *β* is the linear increase in population size per day. Expanding this model to a transmission pair, we assume that all times and parameters are defined along a single forward time axis such that the population size in the donor is simply given by *N_d_(t) = α_d_ + β_d_t*, while the population size in the recipient is given by *N_r_(t) = α_r_ + β_r_(t - t_trans_)*, where subscript *d* indicates the donor and subscript *r* indicates the recipient. The time of transmission is indicated as *t_trans_* when the population size is *N_d_(t_trans_*) in the donor and *α_r_* in the recipient.

**Figure 1.**
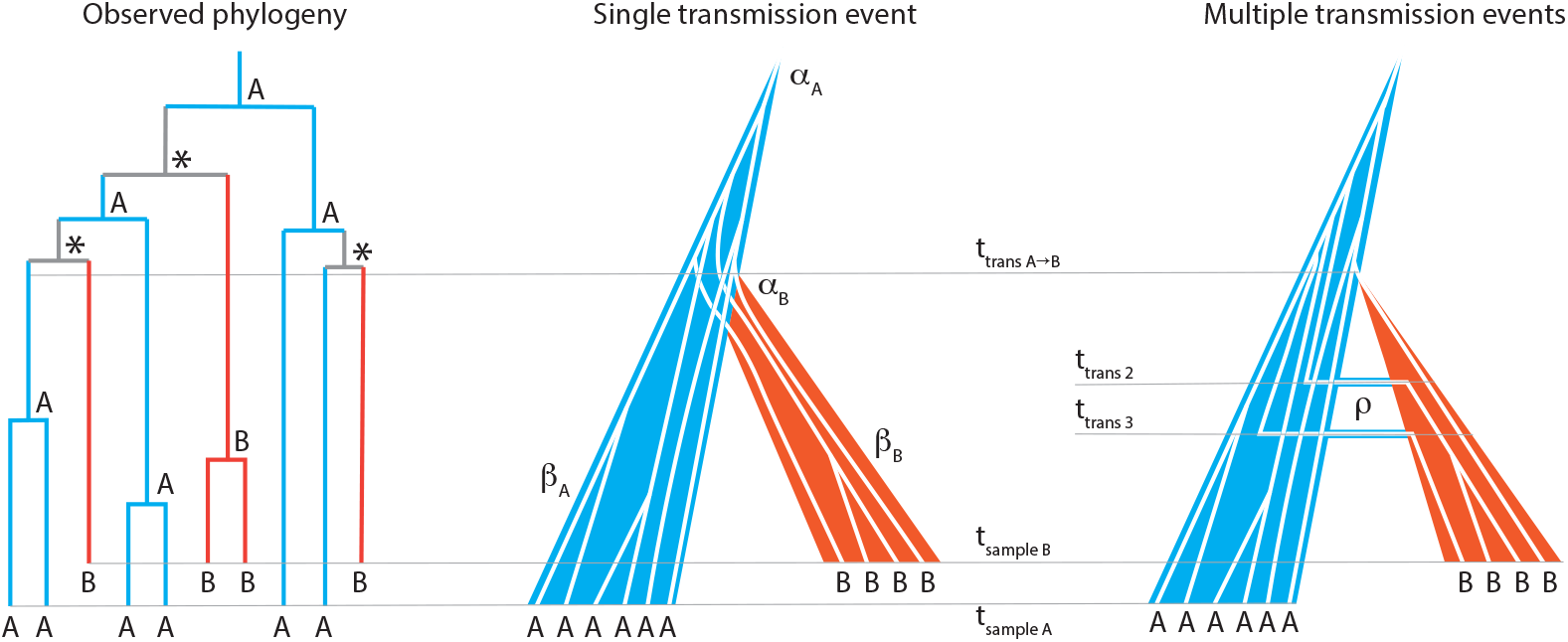
Phylogenetic assessment of transmission scenario. Given a joint donor-recipient HIV-1 phylogenetic tree that suggests transmission of multiple lineages, two transmission scenarios are possible: 1) Transmission of multiple lineages at a single transmission event (co-infection), or 2) transmission of single lineages at multiple events (super-infection). In this example, host A (blue) is donor and B is recipient (red). In the observed phylogeny the root host-label (A, B, or equivocal [*]) is derived by standard maximum parsimony. At time of transmission (ttrans a->b) either multiple lineages are transmitted (single transmission event with aB lineages) or the initial transmission takes place (multiple transmission events each at a=1). Additional transmissions (migration) occurs at later time points (t_trans 2_ and t_trans 3_) at rate *p*. The effective populations grow at β_A_ and β_B_ in donor and recipient, respectively. Samples with individual HIV-1 clonal sequences are taken at t_sample A_ and t_sample B_, respectively.

In the multiple transmission scenario we assume that single virus lineages are passed via sexual contact to the female partner at rate *ρ* and to the male partner at rate *ρ*/2 The half factor corresponds to the reduced rate of female to male transmission (BOILY *et al*. 2009). The population sizes are given by the same equations as in the singular transmission scenario, but where *α_r_* = *α_d_* = 1. We also assume that *ρ* is small enough that *N(t)* is not significantly affected by the migration of lineages between the donor and recipient. We assume that all extant lineages are equally probable to migrate.

### Simulating trees from the joint coalescent model

The derivation of the density of coalescent times for a sample of *k* clones follows Romero-Severson *et al*. (ROMERO-SEVERSON et al. 2014) with modifications to account for the increased rate of coalescence in a population with *k* extant lineages. As before, we define the rate of change in coalescent time as a function of calendar time along the reverse time axis, *s*, as

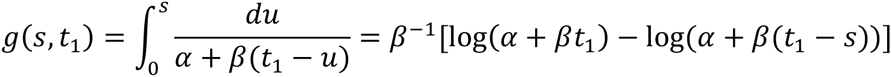

where *t_1_* is the current time. The density of the time to the next coalescent event in Kingman’s n-coalescent with normalized population size for *k* extant lineages is 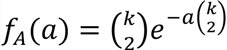 (WAKELEY 2009). Therefore we obtained the density of the time to coalesce in our linear growth model by the transformations

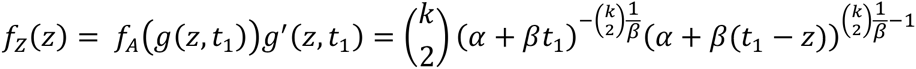

for 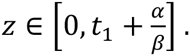

To simulate the time to the next coalescent event we use the inverse cumulative function

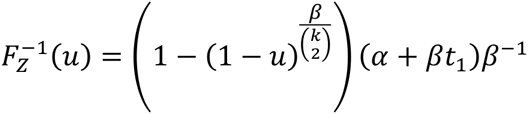

and to simulate the time to the next migration event using the inverse cumulative function

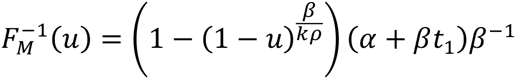

where *u* is a unit uniform random variate. In the singular transmission model a coalescent process was simulated in each of the derived populations of the donor and recipient up to the time of transmission. We define a derived population as a population that only exists in one host after transmission has occurred (in forward time). The derived populations join into a source population at time of transmission, when the lineages from both hosts can freely coalesce (Fig 2). In the source population, a coalescent process was simulated starting with the previous simulations of the derived populations. In the ongoing transmission model four possible events can occur: migration from donor to recipient, migration from recipient to donor, coalescence in donor, and coalescence in the recipient. At a migration event one random lineage moves from one host to the other. Simulations stop when the infection time of the donor is reached along the reverse time axis.

**Figure 2.**
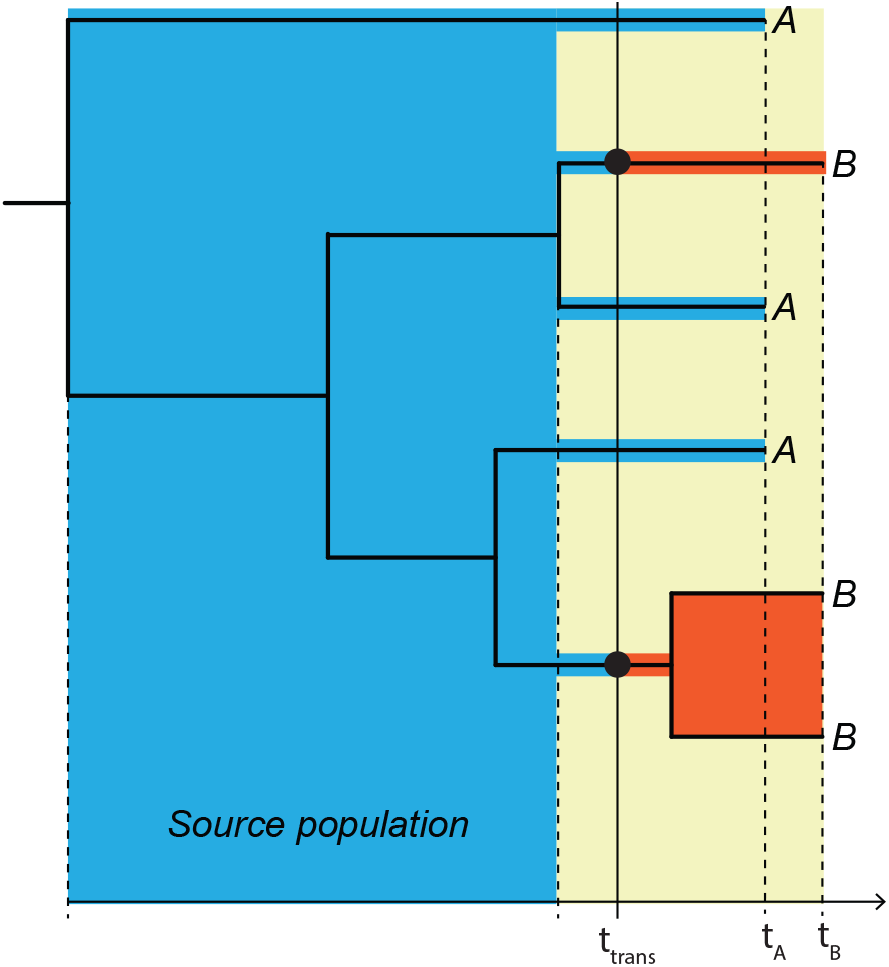
Principle joint donor-recipient time-scaled phylogeny. When a donor (A, blue) infects a recipient (B, red), the possible time-interval when transmission could have occurred (yellow field) is restricted in a time-scaled topology of when the most recent donor-recipient (A-B) coalescence occurred among the sampled lineages and when the recipient was sampled at t_B_. The actual transmission (t_trans_) must have occurred in this interval. The “source population” in direct transmission exists in the donor (blue field), from which at least 2 lineages were transmitted in this example to the donor (red fields). Note that if t_trans_ occurred later at least 3 lineages could have been transmitted.

### Model priors and constraints

To ease interpretation of *α*, we assume that *α* only takes integer values ≥ 1. Because the model is constructed assuming that all parameters and population sizes are continuous it is theoretically possible that the number of lineages that survive though the transmission bottleneck can exceed *α* (e.g. the probability of 5 lineages surviving a bottleneck of size 4 is extremely small but formally non-zero). This incongruity virtually never occurs, however to avoid this situation, we forced a coalescence with branch lengths zero in any case where the number of extant lineages exceeds *N(t)*. We also assume that the donor was infected with a single lineage, *α_d_* = 1.

To constrain the linear growth rate in the recipient, we assumed that the ratio of the population sizes in the donor and recipient is equal to the empirically observed ratio of pairwise diversity between the donor and recipient.

To match the data we wanted to analyze here, we assume that at the time of sampling both donor and recipient were treatment naïve and did not have an AIDS diagnosis. Based on that, and a lack of other relevant epidemiological information, we assumed a uniform distribution of infection times from 0 to 12 years. We assume that the population growth rate in the donor is drawn from β*_d_*~Exponential(15^−1^) units per day. This distribution includes growth rates that correspond to most of the published estimates of the HIV within-host effective population numbers (LEIGH BROWN 1997; NIJHUIS *et al*. 1998; PENNINGS *et al*. 2014). In the case of a singular transmission event we assume that the donor transmits on average 0.7% (s.d. 0.9%) of the current effective population number (Beta(0.5,70) distributed) to the recipient. In the ongoing transmission case we assume the transmission rate from the male to female partner is a uniform random variable between 0 and 2 per day, *ρ*~Uniform(0,2).

### Phylogenetic measures for approximate Bayesian computation

For a tree with taxa from two hosts, “A” and “B”, we used the following statistics to define the probability that a simulation should be accepted: the root label, the topological class, the number of monophyletic clades of one of the host labels, the total number of substitutions in the tree, and the average pairwise distance between pairs of taxa with mismatched host labels. The root label is defined as the maximum parsimony host assignment of the root (“A”, “B”, or ambiguous). The topological relationship can be one out of three classes: MM (both host sets of taxa are monophyletic), PM (taxa from one host forms a monophyletic clade that inserts into the sample of the other host forming a paraphyletic clade), and PP (taxa from one host are paraphyletic to the other host’s taxa that are polyphyletic, or both host’s taxa are polyphyletic). Root label and topological class have been demonstrated to be associated with the epidemiologic relationship between two sampled hosts (ROMERO-SEVERSON *et al*. 2016). The number of monophyletic clades of the putative recipient in the joint tree defines the minimum number of transmitted lineages. Note that it probabilistically informs the number of transmitted lineages, e.g. a large number of transmitted lineages is generally—but not always—inconsistent with an observed single monophyletic clade. With the root label assigned to “A”, the number of “B” clades in the sample is counted by applying Dollo’s law (DOLLO 1893), which logically follows from the irreversible fact that the donor was infected before the recipient. In principle, this translates on the tree to first assigning the “A” label to each node on a root to “A”-tip path, and then counting the minimum “A” to “B” transformations needed to observe the tip labels. We call each such “B” lineage a “drmonophyletic clade”, including clades with only one “B” taxon.

In the single transmission of multiple lineages scenario, rescaling the tree using a molecular clock identifies the time interval during which transmission could have occurred. In that case the first coalescence going towards the root between a “A” and “B” lineage defines the time of when the tree describes the HIV-1 evolution in the donor, i.e., the “source population” (ROMERO-SEVERSON *et al*. 2016). Thus, the time during which transmission could have occurred spans from the time of the sampling of the recipient back until the time that defines the source population (Fig 2). The total number of substitutions is calculated by assuming a Gamma distributed uncorrelated relaxed molecular clock with a mean evolutionary rate at 6.7 (s.d. 4.2) ×10^-3^ substitutions/site per year that informs both the infection time and the within-host population growth rate (LEITNER AND ALBERT 1999). Finally, the mean number of substitutions between donor-recipient pairs informs the donor-recipient transmission time and is linked to the number of transmitted lineages. Note that the possible transmission time interval in the multiple transmissions of single lineages scenario is undefined because it does not have a strict logical boundary going towards the root. Hence, we do not use that model to estimate the time of the original transmission.

### Sampling from the posterior density of parameters

To account for variance in the phylogenetic trees that are consistent with the sequence data, we calculated the statistics described above for each tree and normalized the results to obtain distributions of the statistics conditional on the data and phylogenetic model. We considered two separate probabilistic sampling schemes based on either the topological statistical alone—without having to specify the evolutionary rate—or the full set of tree statistics. Both schemes consider each statistic as an independent test that is probabilistically passed with the empirical probability of the statistic. For example, if the empirical probability of the PP topology is 1.0 (every tree in the posterior had a PP topology) then any simulation that does not produce a PP tree is rejected. Likewise, if the probability of the “A” root label is 0.93, then a simulation with root label “A” will be accepted 93% of the time. Each statistic is considered independent such that the total probability of accepting a parameter is proportional to the product of each of the simulated statistics.

The first sampling scheme is based only on the topological class and root label statistics, while the second sampling scheme is based on the topology, root label, number of monophyletic clades in “A”, the total number of substitutions in the joint tree, and the average distance (in substitutions) between “A”-“B” pairs.

We sampled 10^7^ parameter sets from the prior, for each of the 4 possible models (“A” donor, singular transmission; “A” donor, multiple transmissions; “B” donor, singular transmission; and “B” donor, multiple transmissions). We considered the ratio of the number of accepted parameters as an approximation of a Bayes factor for the model. We took the marginal means of the accepted samples to be the point estimate for the model parameters and the appropriate quantiles to define the 95% credible intervals.

### Study subjects

To test our model and ABC framework, we analyzed a set of sequences from three HIV-1 infected subjects (MP1, MP2, MP3) that had been part of a forensic HIV transmission investigation, where male MP2 had accused female MP1 of intentional transmission, and where MP2 infected MP3, a later female sexual partner. MP1 and MP2 subjects had a history of intravenous drug use, but MP3 did not. Thus, based on the epidemiological record, MP1 and MP2 could potentially have infected each other via either sexual contact or needle injection, but transmission between MP2 and MP3 could only have been through sexual interaction. Based on maximum likelihood (ML) phylogenetic reconstruction of HIV-1 env DNA sequences, MP1 taxa were separated from MP2 taxa by multiple local control and database sequences (Fig S1). Hence, MP1 was highly unlikely to have infected MP2 or MP3. However, the phylogenetic reconstruction was consistent with HIV-1 transmission from MP2 to MP3. The criminal investigation concluded that MP1 had not infected MP2, in part based on the phylogenetic evidence (Fig S1). That investigation used the case sequences in this paper plus 119 env sequences selected from Portuguese and publicly available databases. In the general framework above, MP2 can been seen as host “A” and MP3 as host “B”.

### DNA sequencing

Chromosomal DNA was extracted from infected PBMC’s of each subject using Wizard Genomic DNA Purification Kit (Promega) according to the manufacturer recommendations. Nested PCR was done to obtain a 534 bp fragment from the C2V3 env region (HXB2 positions 6858-7392). Thermal cycling conditions were as previously described (BARTOLO *et al*. 2009). PCR products were cloned into the pCR™4-TOPO vector (Invitrogen), using the TOPO TA Cloning Kit (Invitrogen) according to the manufacturer’s instructions. DNA sequencing was performed using the BigDye Terminator V3.1 Cycle sequencing Kit (Applied Biosystems) and an automated sequencer (3100-Avant Genetic Analyzer; Applied Biosystems, Foster City, CA). We derived 31, 20, and 19 sequences from MP1, MP2, and MP3, respectively.

### Phylogenetic reconstruction

HIV-1 sequences were aligned using MAFFT with the L-INS-i algorithm (KATOH AND TOH 2008). Maximum likelihood phylogenetic trees were inferred using PhyML (GUINDON *et al*. 2005) under a GTR+I+G substitution model, 4 categories Gamma optimization, with a Bio-NJ starting tree and best of NNI and SPR search, and aLRT SH-like branch support (ANISIMOVA AND GASCUEL 2006). The posterior distribution of trees was sampled using MrBayes (RONQUIST AND HUELSENBECK 2003) under the same model parameterization as the PhyML trees. Two Markov chains were run for 20 million steps each. Removing 25% of the chain as burn-in, combining the chains, and sampling every thousandth tree, we obtained 30,000 independent trees from the posterior distribution of trees.

### Data availability

Sequences have been deposited in GenBank under accession numbers KT123041-KT123171.

## RESULTS

### Direct transmission of multiple, diverse phylogenetic lineages

Using the MP1 population as outgroup, the inferred rooted ML tree suggested that MP3 was infected with at least 7 independent phylogenetic lineages from MP2 (Fig 3). The ML tree thus indicated a paraphyletic-polyphyletic (PP) topological relationship (Romero-Severson *et al*. 2016), strongly suggesting direct transmission from MP2 to MP3.

**Figure 3.**
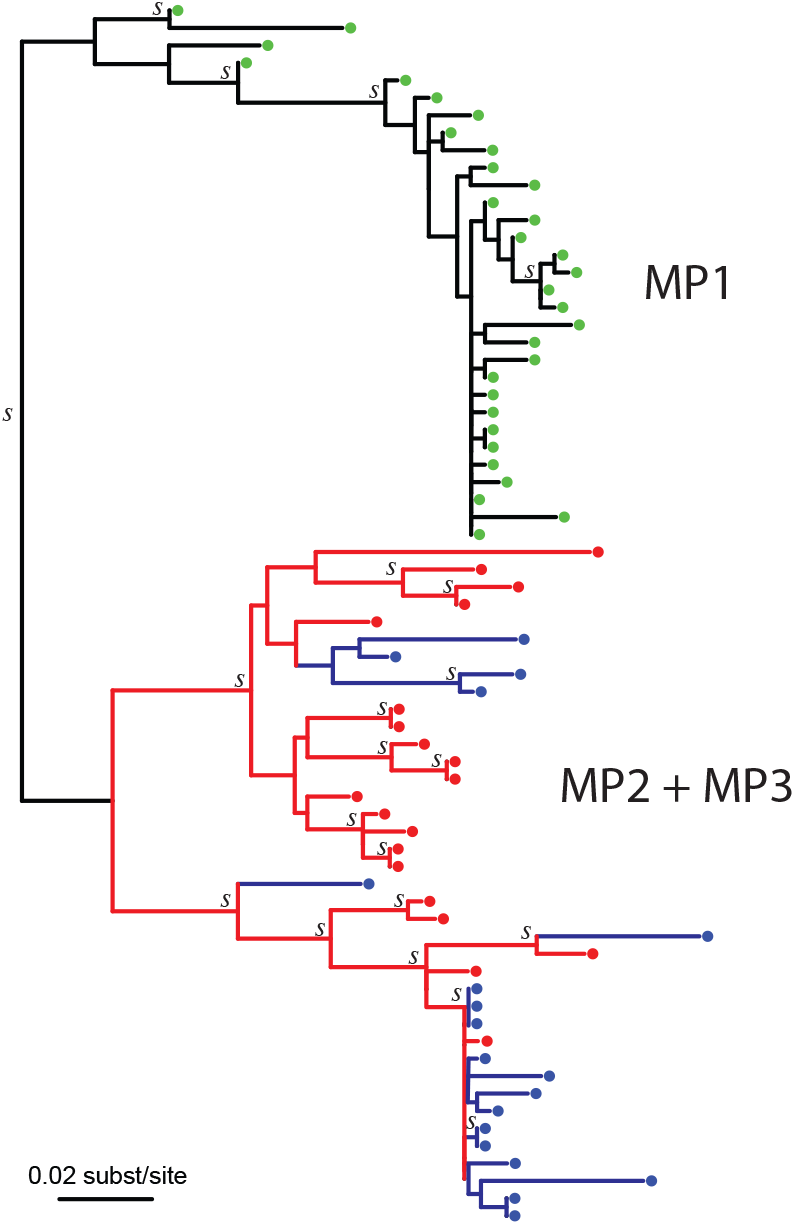
Maximum likelihood reconstruction of the MP2-MP3 joint HIV-1 env phylogeny. MP1 (yellow) did not infect either MP2 or MP3 (Fig S1), and is used to root the MP2 (red) and MP3 (blue) HIV-1 tree. Clades with aLTR support (>0.90) are indicated with a “S”. The topology of this tree suggested that at least 7 lineages were transmitted from MP2 to MP3. Because the branch lengths were zero or near zero in the bottom clade, we added a small distance for readability purpose to show the 4 possible transmitted lineages that the topology suggested in this clade. Partially to avoid depending on this single (best) tree, we evaluated 30,000 posterior trees presented in in Figure 4.

However, the ML tree does not give any sense of the variance in the topological signal that suggests that MP2 is the likely donor. In the posterior distribution of trees (obtained using MrBayes) we found that 93% of the trees had a PP topology with an MP2 root label. Likewise, the mean number of monophyletic clades in the posterior (7.7) was close to the number of monophyletic clades in the ML tree (7), however, the range in the posterior was quite large (4-14, Fig 4A). Note that the bottom 4 monophyletic clades in the ML tree (Fig 3) are only very weakly separated considering branch lengths, and thus it is no surprise that the posterior distribution of trees display a range of possible values. It is important to point out that the number of monophyletic clades is the *minimum* number of lineages establishing an infection; the actual number of infecting lineages can be much higher due to extinction of founding clades and suboptimal sampling of extant lineages in the donor and recipient.

**Figure 4.**
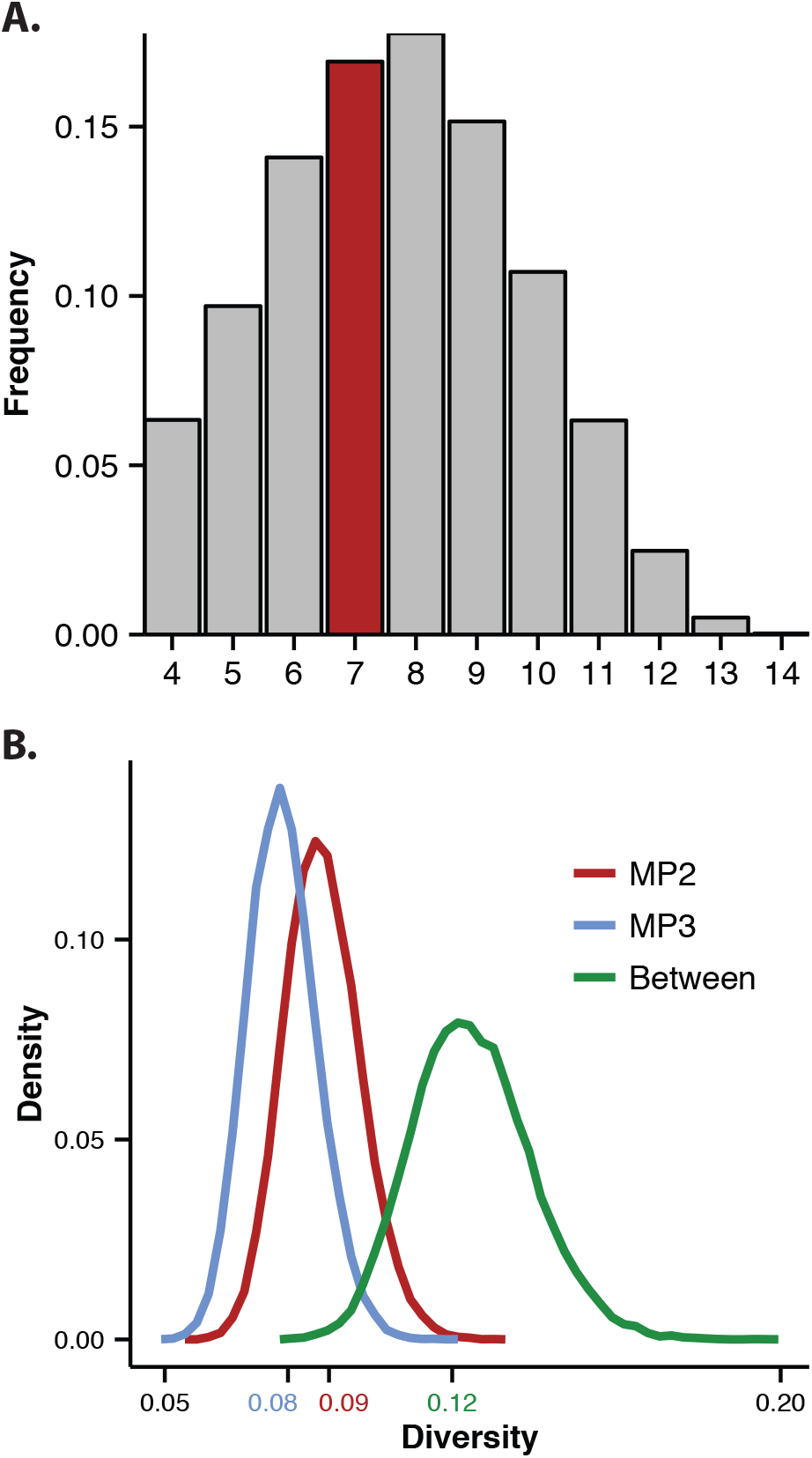
Evaluation of posterior tree sample. 30,000 MrBayes trees were sampled after burn-in to evaluate the possible range of inferred minimum number of transmitted lineages and sampled diversity in the hosts. **(A)** The number of MP3 monophyletic clades in the joint MP2+MP3 trees ranged from 4—14 in the Bayesian posterior MCMC sample with a mean of 7.7 near the ML estimate at 7. **(B)** The diversity as measured by the patristic tree distance (substitutions/site) in hosts MP2, MP3, and between the HIV-1 populations.

The high diversity amongst the taxa from MP2 and MP3 also supports the idea of a high degree of shared diversity between MP2 and MP3 (Fig 4B). Comparing the within-host diversity at times of sampling showed that MP3 had only a little less diversity than MP2 (the mean pairwise taxa distance in MP2 and MP3 was 0.088 and 0.079 substitutions/site, respectively, again in agreement with the ML tree at 0.090 and 0.075 substitutions/site, respectively). Furthermore, the between MP2 and MP3 population distance was somewhat larger (0.124 and 0.122 substitutions/site for posterior tree sample and ML tree, respectively). Together, these distances indicated that a large amount of diversity indeed had been transmitted.

The transmission of multiple lineages and the large diversity can only occur if there was either a large diverse population transmitted from the donor or if there were multiple transmission events between MP2 and MP3.

### Evidence for direction and frequency of transmission

To evaluate how so many lineages and so much diversity was transmitted, we considered 4 possible scenarios to explain the data: MP2 or MP3 as the donor, and transmission at either a singular event or at multiple occasions (Fig 1). We considered the ratio of acceptance rates of samples from the priors based on either the topological statistics or the full set of tree statistics in the 4 possible scenarios as the relative evidence of one hypothesis over the other.

The evidence based on the full tree statistics favored MP2 as the donor in both the single and multiple transmission scenarios; Bayes factor (BF) was 24 for the singular transmission case, 9.9 for multiple transmissions, and 12 for combined. If only topological statistics were used, the evidence was weaker but still favored MP2 as the donor (BF 3.6 in singular transmission, 2.5 in multiple transmissions, and 2.8 when combined). Thus, regardless of whether transmission occurred once (with multiple lineages) or many times (with one lineage at each time), the evidence points to MP2 as the donor.

Both the topological and full tree statistic very weakly favored the ongoing transmission case (BF 1.3 and 2.1, respectively). From a purely statistical point of view, we could establish MP2 as the donor, but it was unclear whether there had been a single or multiple transmission events between the MP2-MP3 pair.

### Point estimates and credible intervals of model parameters

To further analyze support for one transmission scenario over the other, we evaluated the coalescent model parameters (Tab 1). With priors informed by available clinical and epidemiological information, the infection time of MP2 and linear growth rate β in MP2 were robust to our singular or multiple transmission event hypotheses. Infection time of MP3 and linear growth rate in MP3 were not, however. Because β_MP3_ is a function of the infection time of MP3, the lack of robustness in β_MP3_ is due to the lack of identifiability of the infection time in the ongoing transmission case. Similarly, the initial transmission time in the multiple transmission scenario is not well identified (posterior is close to prior). This is likely due to the fact that one cannot identify where the process starts (the infection time) if the rate of additional transmissions (migration) is not well constrained.

### Extreme transmission rate makes multiple transmission events implausible

The level of ongoing transmissions that is consistent with the data is very high (ρ in Tab 1 & Fig 5). In our model MP2 and MP3 reinfect each other at rate *ρ* and *ρ*/2 respectively over the time period from when MP3 was first infected until sampled (corresponding to the yellow area in Fig 2). The mean posterior (*ρ*= 1.3) indicates that transmission events occur more than once per day, which is implausible as it implies very high contact rates and greater than ever reported risk of HIV-1 transmission per heterosexual contact (BOILY *et al*. 2009). Even the lower posterior bound (*ρ*= 0.3) is almost certainly biologically implausible.

**Figure 5.**
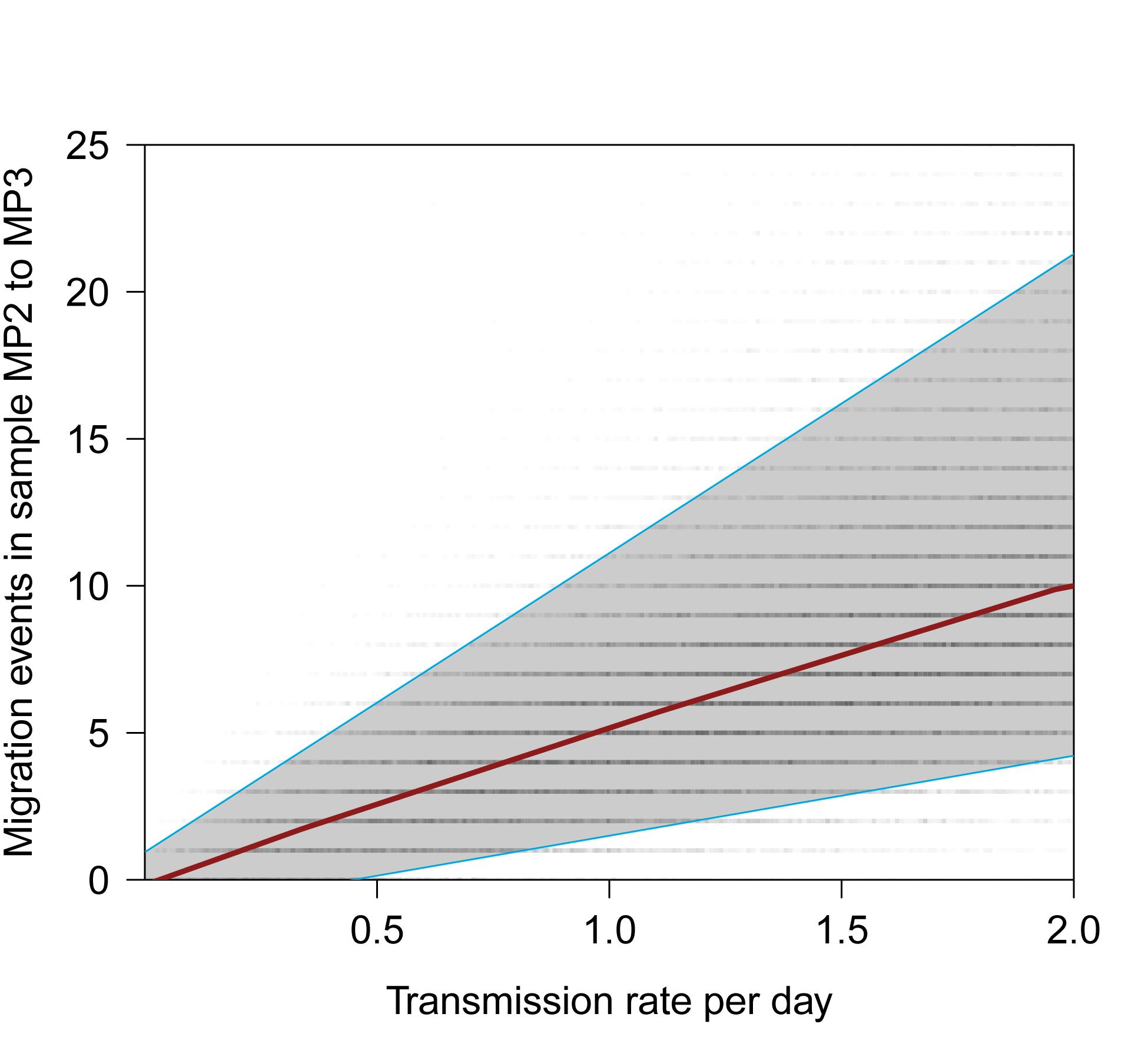
Migration analysis in the multiple transmission events scenario. Simulation results using our ABC coalescent-based method showed that the transmission rate ρ (x-axis) implied a very high number of migration events in the modeled population to explain the number of MP3 monophyletic clades observed in the sample (y-axis). Red line, median trend of 50,000 simulations; grey envelope with blue edges, 5-95% interval; grey open circles, individual simulation results.

Thus, while the ongoing transmission model explains the data well, the model only works at very high levels of transmission between MP2 and MP3 (Fig 5). This is due to the fact that the number of sampled lineages (20 sequences from MP2 plus 19 sequences from MP3) is much smaller than the population sizes in MP2 and MP3. To observe any transmission events in the very small subset of the population that was sampled, there must be a very high rate of ongoing transmission between MP2 and MP3 (Fig 5). Therefore, from an epidemiological point of view, the implausible posterior transmission rate that is needed to support our data under the ongoing transmission scenario rather lends support for single transmission of multiple lineages.

### Robust estimation of transmitted diversity

Effective population size (Ne) is a model construct based on an idealized population and is thus an abstract formalism that can be difficult to interpret. However, Ne can be linked to the more concrete and measurable population diversity. our maximum posterior estimate of *α* = 116 does not necessarily imply that exactly 116 virions were transmitted; we can, however, estimate the diversity of the establishing inoculum by simulating an evolutionary process from the posterior distribution of parameters corresponding to the transmitted lineages. Hence, while we cannot say exactly how many physical virions established the infection, we found that the mean diversity of the establishing population after transmission was robust over the posterior distribution of parameters. Notably, the diversity of the transmitted population was nearly as diverse (0.069 substitutions/site) as was later observed among the sequence clones in MP2 and MP3 at time of sampling (see 1^st^ results section). Note also that the total diversity at time of sampling was likely larger than that observed among the clones.

To investigate the effect of α on mean diversity at transmission, we first measured diversity as α increased while other parameters were fixed at their maximum posterior values (Tab 1). As the number of transmitted lineages increases linearly the expected transmitted diversity initially increases rapidly (m1 in Fig 6). Allowing all model parameters to vary, we see a similar increase in diversity with increasing α (m2 in Fig 6). In both situations diversity of the founding infection does not increase beyond α>20. Note that the diversity plateaus at a higher level when only a is varied; when all remaining parameters are sampled from the posterior they have a compensating effect (e.g. lower *β_MP2_*) to explain the robust diversity estimate. Naturally, as higher α increases the certainty that transmission involves high diversity, a corresponding decrease in the standard deviation is observed (Fig 6). Thus, while there is a non-linear relationship between α and transmitted diversity, we can be reasonably certain that MP2 infected MP3 with a highly diverse inoculum. This result is robust even if we consider only the range of the absolute minimum number of transmitted viruses implied by the posterior distribution of the number of monophyletic clades in MP3 (4-14).

**Figure 6.**
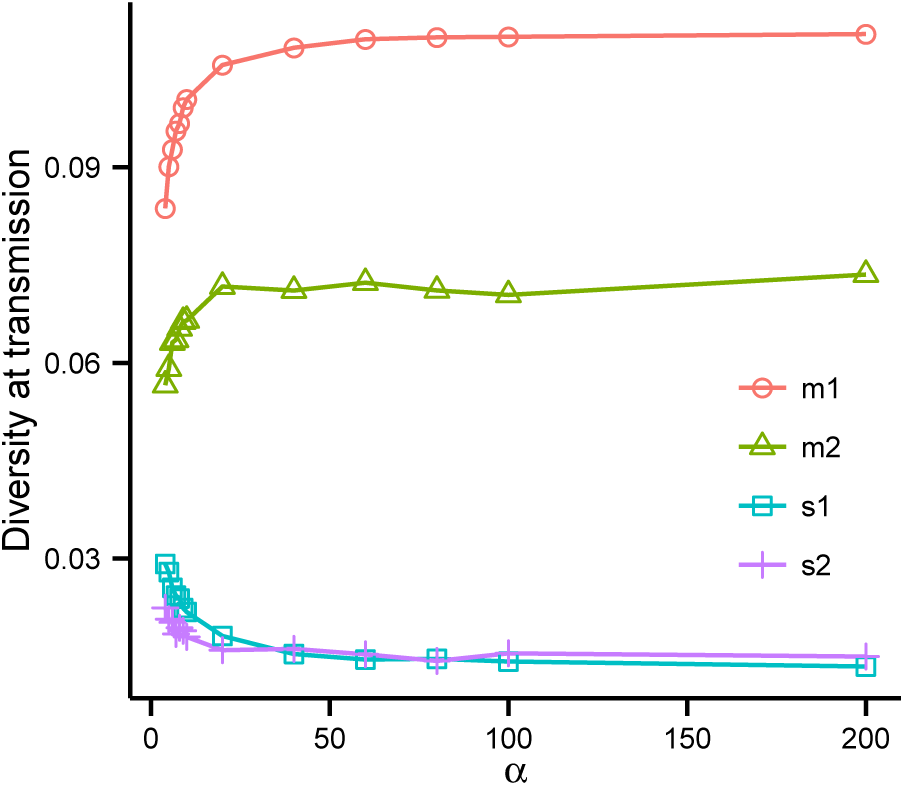
Inference of the transmitted diversity among lineages. Simulation m1 shows how the transmitted diversity changes as only a increases (with all other model parameters fixed at the maximum posterior values in Tab 1). Simulation m2 shows the same trend when all model parameters are optimized at different a. s1 and s2 are the corresponding standard deviations.

## DISCUSSION

In this study we show how to apply previously described theoretical evaluations of epidemiological linkage to a real HIV-1 transmission case that involved a highly diverse founding HIV-1 population. We show that one can simultaneously estimate direction, diversity, and frequency of the transmission event(s). We used a previously developed within-host coalescent framework (ROMERO-SEVERSON *et al*. 2014), and expanded it by allowing additional transmission events (migration) between the hosts. Inference was achieved using an ABC method informed by topological and distance-based tree statistics, which allowed Bayes factor comparisons between alternative epidemiological scenarios.

The transmission between MP2 and MP3 involved many lineages, probably more than we could observe among the limited sample of HIV-1 sequences derived from the patients. It is impossible to know exactly how many lineages were transmitted with these data. Our ABC framework can however estimate the diversity that was transmitted, and arguably this measure is more important from a clinical perspective as it may relate to how difficult it is to combat the incoming virus for the immune system, antiviral drugs, and future vaccines. We estimated that the inoculum that infected MP3 had a diversity of 0.069 substitutions/site, which is very high (corresponding to years of diversification). This level of diversity is equivalent to an incoming effective population size, α, of either ≈100 or alternatively there was additional transmission events between MP2 and MP3 at levels that are highly unrealistic compared to empirical estimates of heterosexual transmission rates (BOILY *et al*. 2009). Thus, the transmission of the degree of diversity in this case seems to be the result of a single transmission with a very diverse inoculum involving many phylogenetic lineages from the donor.

HIV-1 co-infection has been defined as infection of several HIV-1 genetically diverse virions before seroconversion (typically 21 days after infection (COHEN *et al*. 2011)) or within a somewhat later time (3-6 months) when a strong immune response has developed to the initial inoculum, and super-infection as additional infections after the strong immune response has been established (VAN DER KUYL AND CORNELISSEN 2007; RONEN *et al*. 2013). In addition, super-infection is often thought of as an additional infection from another donor than the initial one. In the transmission case we studied here, both co- and super-infection was evaluated involving only the original donor and recipient, a stable heterosexual couple. Thus, with repeated contacts over time, transmissions may span and blur the defined periods of co- and super-infection. Importantly, HIV-1 evolves significantly during any period >1 month (SKAR *et al*. 2011), putting later transmitted variants somewhere in between the genetic diversity possible from co-infection from the original donor and superinfection from another donor, further blurring the co- and super-infection distinctions. Thus, while super-infection involving multiple donors appears rare (VAN DER KUYL AND CORNELISSEN 2007), given the fact that 20-40% of sexual infections involve >1 genetic variant (KEELE *et al*. 2008; SALAZAR-GONZALEZ *et al*. 2009; LI *et al*. 2010; Rieder *et al*. 2011), ongoing transmission between stable couples as investigated here may be more common than previously realized. On the other hand, at least in the case we studied here, ongoing transmission seemed unrealistic as it implied an impracticable high transmission rate (BOILY *et al*. 2009). Our study provides the first results of modeling single versus ongoing transmission events to explain how multiple lineages could end up in the recipient. A possible extension to our framework could be to allow for transmission of >1 lineage at multiple times, but without additional data, e.g., frequent longitudinal and deep sampling, there would not be enough power to identify how many variants that were transmitted at each possible occasion.

We have recently shown that a joint HIV-1 phylogeny of two epidemiologically linked hosts may reveal direction and directness when >1 lineage has been transmitted (ROMERO-SEVERSON *et al*. 2016). As predicted by our topological evaluation, the epidemiological record confirmed that the inferred paraphyletic-polyphyletic (PP) tree resulting from the MP2-MP3 transmission identified MP2 as donor with no intermediary link to MP3. We note that the MP2 donor assignment was independent of whether transmission occurred once with multiple lineages or many times with single lineages. We also expanded the theoretical predictions with evaluations of many possible trees that could reasonably explain the sequence data, i. e., by evaluating a posterior tree sample derived from a Bayesian MCMC tree search using MrBayes. We show that in our case a ML tree reconstructed with PhyML (using NNI+SPR search) gives a good point estimate of the number of transmitted lineages in the sample, but that this gives no idea of the possible range. We hypothesize that there are situations when a ML-based estimate may not agree with the maximum posterior estimate. This becomes especially true in more complex models when parameters can compensate for each other to explain the data.

In conclusion, taking phylogenetic uncertainty into account, we have created a framework that can evaluate how much diversity is transmitted, and whether transmission occurs once or over a period of time. We argue that estimating the transmitted diversity in the inoculum may reveal more about how difficult a transmission would be to prevent or fight than trying to find the exact number of transmitted lineages.

## ACKNOWLEDGEMENTS

Research reported in this publication was supported by the NIAID/NIH under award number R01AI087520 and by grants PTDC/SAU-EPI/122400/2010, VIH/SAU/0029/2011 and UID/Multi/04413/2013 from Fundação para a Ciência e Tecnologia (FCT), Portugal. Inês Bártolo was supported by a post-doc fellowship (SFRH/BPD/76225/2011) from FCT, Portugal. Ingo Bulla was supported by a postdoc fellowship (BU 2685/4-1) from the Deutsche Forschungsgemeinschaft.

---

**Fig S1.**
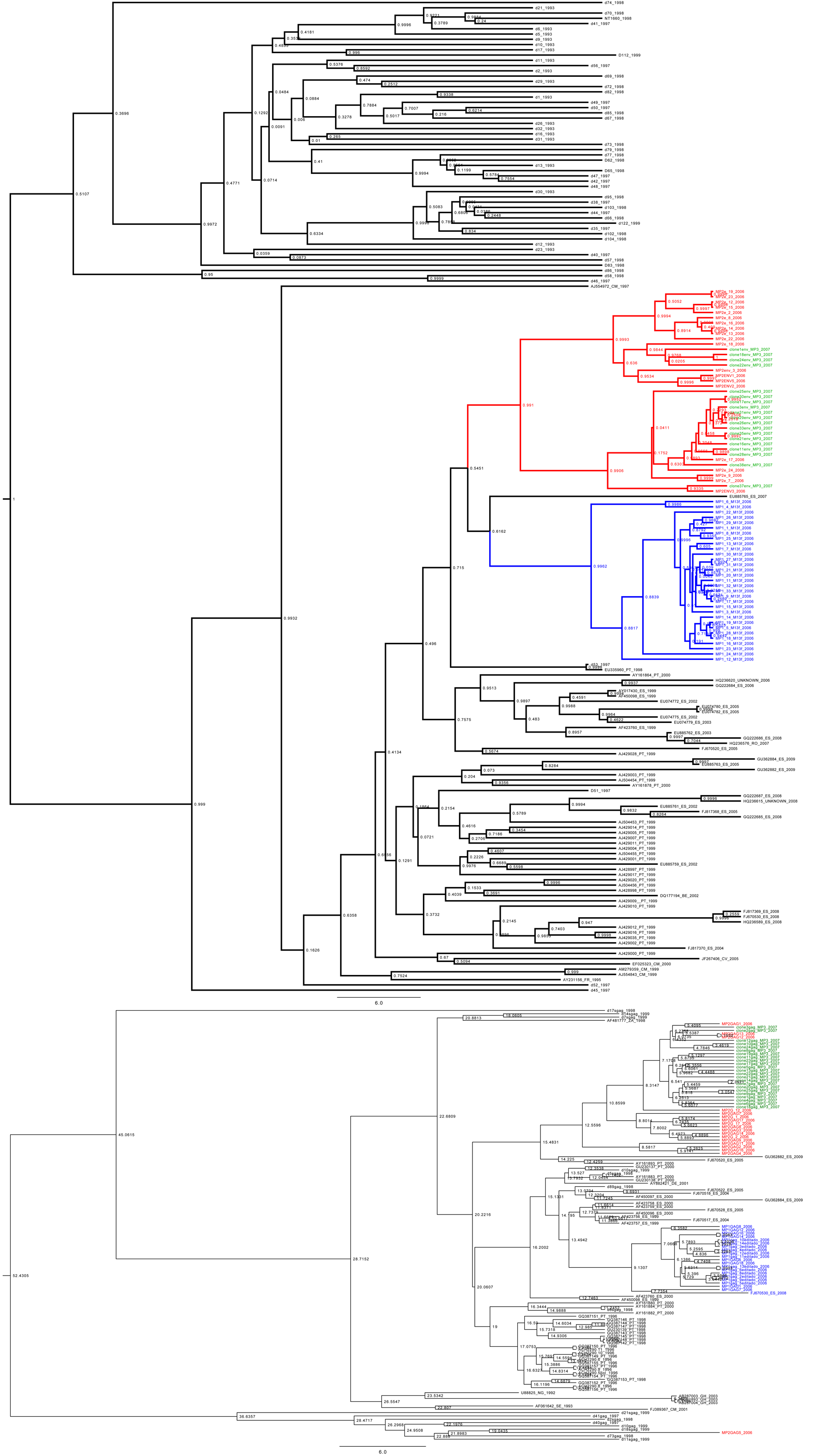
Time-scaled phylogeny of MP1, MP2, and MP3 HIV-1 populations compared to local and database control sequences. MP1, MP2, and MP3 sequences are indicated by color, and control sequences in black. Numbers at nodes indicate posterior support values.

